# Increasing Grain Yield in Bread Wheat (*Triticum aestivum* L.) by Selection for High Spike Fertility Index

**DOI:** 10.1101/2020.10.30.361782

**Authors:** Ana Clara Pontaroli, María Pía Alonso, Nadia Estefania Mirabella, Juan Sebastián Panelo, María Fiorella Franco, Leonardo Sebastián Vanzetti, Máximo Lorenzo

## Abstract

Spike fertility index (SF) has been proposed as a promising selection criterion for increasing grain yield (GY) in bread wheat. Here, changes in GY and related traits after simulated selection (10% intensity) for high SF or high GY *per se* were assessed in two RIL populations: (I) Avalon/Glupro and (II) Baguette 10/Klein Chajá, and in (III) advanced lines from a breeding program. Grain yield, SF, grain number per unit area (GN), grain weight (GW), test weight (TW) and grain protein content (GPC) were determined. Regardless of the environmental conditions, simulated selection for high SF always resulted in GN increases (between 1.6 and 27.4%). Average GY increase observed after selection for high SF (11.5%; N=10; SEM=5.8) did not differ (p=0.92) from the average GY increase observed after selection for GY *per se* (11.8%; N=10; SEM=4.9). Grain weight, GPC and TW tended to decrease with selection for high SF; however, these trade-offs might be avoided by concurrent selection. Our findings validate the use of SF as a selection criterion for increasing grain yield in bread wheat.

## INTRODUCTION

Grain yield is one of the key targets of bread wheat (*Triticum aestivum* L.) breeding programs worldwide. Under a scenario of increased global demand for grains and limited possibilities of expanding cropping areas, breeding efforts must focus on raising annual yield increase rates (CIMMYT, 2019). One way to achieve this is by using yield-related, high-throughput phenotypic variables as selection criteria.

Grain yield in wheat is determined mainly by the number of grains per unit area (GN) and, to a lesser extent, by grain weight (GW). Therefore, variations in GN best explain yield variation (Abbate et al., 1998; Borrás et al., 2004; Elía et al., 2016; Ferrante et al., 2017; Lynch et al., 2017; Shearman et al., 2005; Slafer et al., 1990). However, this is a difficult trait to select for in early generations of a breeding program, in which not enough seed is available. Fischer (1984) proposed that, under non-limiting growing conditions (i.e., without water or nutrient limitations and in absence of pests and diseases), GN is determined by the product of (i) the duration of rapid spike growth period, (ii) crop growth rate during the spike growth period, (iii) dry weight partitioning to spikes during the spike growth period, and (iv) the number of grains per unit of spike dry weight, i.e., a “spike fertility” index (SF) (Abbate et al., 1998) or “fruiting efficiency” (Ferrante et al., 2012). The spike dry weight at anthesis is a complex trait to measure; thus, chaff dry weight at maturity has been proposed as its surrogate (Abbate et al., 2013). Then, the spike fertility index at maturity [also termed fruiting efficiency at maturity; Fischer and Rebetzke (2018)] is calculated as the number of grains produced per unit of chaff dry weight.

There is extensive evidence in the literature that GN is tightly linked to SF (Abbate et al., 1998; Acreche et al., 2008; Alonso et al., 2018b; Foulkes et al., 2015; Shearman et al., 2005; Terrile et al., 2017). In the reference method (9), SF determination is carried out at anthesis, but this is destructive and very time consuming. A high throughput, non-destructive method for assessing the trait in small samples at maturity has then been proposed (Abbate et al., 2013). In addition, SF at maturity has been shown to have high heritability, substantial genetic variability and low genetic by environment interaction (Alonso et al., 2018b; Martino et al., 2015; Mirabella et al., 2016). Because of all these features, SF has been regarded as a promising selection criterion for increasing GN (and yield) in breeding programs (Abbate et al., 2013; Elía et al., 2016; Fischer, 2007, 2011; Fischer & Rebetzke, 2018; Foulkes et al., 2010; García et al., 2014; González et al., 2011; Lázaro & Abbate, 2012; Martino et al., 2015; Mirabella et al., 2016; Slafer et al., 2015; Terrile et al., 2017). As a matter of fact, Alonso et al. (2018b) have shown that selection for high SF, either solely or in combination with selection for high grain yield, effectively increases yield, resulting in higher and more stable yields than if selecting for high yield *per se*. These encouraging results, obtained with a single RIL population under non-limiting environmental conditions, need further validation before such selection strategies can be implemented in a breeding program. Also, the assessment of possible trade-offs that could arise as a byproduct of these schemes should be carried out. Therefore, the aim of this work was to determine the effectiveness of selection for high SF in raising grain yield and its effect on other variables, by carrying out simulated selection and analyzing the response to this selection in a combination of diverse genetic materials (RILs and advanced lines) and an array of environmental conditions.

## MATERIALS AND METHODS

### Field experiments and plant material

Field experiments were carried out at the Balcarce (37°45’ S; 58°18’ W; 130 m a.s.l.) and Marcos Juárez (32°43’S; 62°06’W; 112 m a.s.l.) Agricultural Experimental Stations of the Instituto Nacional de Tecnología Agropecuaria (INTA), Argentina. Characteristics of experiments are summarized in Table 1.

**Table 1.** Description of field experiments carried out in this study, as organized in Sets of experiments (exps.) I to III. Genetic material included in each set, N (number of individuals), exp. name, location, year, exp. design (RCBD=randomized complete block design; ARCBD=augmented randomized complete block design), No. reps. (number of replications), growing conditions, plot dimensions and data collected (SF=spike fertility index; GY=grain yield; GN=grain number per unit area; GW=grain weight; TW=test weight; GPC=grain protein content). Data from exps. BFE13, BFE14 and BFE15 have been partially published by Alonso et al. (2018a;b).

| Set of exps. | Genetic material | N | Exp. name | Location | Year | Sowing date | Exp. design | No. reps. | Growing conditions | Plot dimensions |  |  |  | Data collected |  |  |  |  |  |
| --- | --- | --- | --- | --- | --- | --- | --- | --- | --- | --- | --- | --- | --- | --- | --- | --- | --- | --- | --- |
|  |  |  |  |  |  |  |  |  |  | Area (m<sup>2</sup>) | No. rows | Row length (m) | Inter-row spacing (m) | SF | GY | GN | GW | TW | GPC |
| I | RIL population (Avalon/Glupro) | 90 | BAG15 | Balcarce | 2015 | June 19<sup>th</sup> | RCBD | 2 | Rainfed | 3.5 | 3 | 5 | 0.2 |  | X | X | X | X | X |
|  |  | 90 | BAG16 | Balcarce | 2016 | July 14<sup>th</sup> | RCBD | 2 | Rainfed | 3.5 | 3 | 5 | 0.2 | X | X | X | X | X | X |
| II | RIL population (Baguette 10/ Klein Chajá) | 146 | BFE13 | Balcarce | 2013 | June 23<sup>rd</sup> | RCBD | 2 | Irrigated | 3.5 | 3 | 5 | 0.2 | X | X | X | X | X |  |
|  |  | 146 | BFE14 | Balcarce | 2014 | July 25<sup>th</sup> | RCBD | 2 | Irrigated | 7 | 7 | 5 | 0.2 | X | X | X | X | X |  |
|  |  | 146 | BFE15 | Balcarce | 2015 | July 15<sup>th</sup> | RCBD | 2 | Irrigated | 7 | 7 | 5 | 0.2 | X | X | X | X | X |  |
|  |  | 146 | MJFE17 | Marcos Juárez | 2017 | June 15<sup>th</sup> | ARCBD | 1 | Rainfed | 0.9 | 3 | 1 | 0.3 | X |  |  |  |  |  |
| III | Advanced breeding lines | 28 | BLE16 | Balcarce | 2016 | August 8<sup>th</sup> | ARCBD | 1 | Rainfed | 7 | 7 | 5 | 0.2 | X | X | X |  |  |  |
|  |  | 28 | BLE17 | Balcarce | 2017 | June 14<sup>th</sup> | ARCBD | 1 | Irrigated | 7 | 7 | 5 | 0.2 | X | X | X |  |  |  |
|  |  | 36 | BLP17 | Balcarce | 2017 | August 6<sup>th</sup> | ARCBD | 1 | Rainfed | 7 | 7 | 5 | 0.2 | X | X | X | X |  |  |
|  |  | 36 | BLE18 | Balcarce | 2018 | June 11<sup>th</sup> | RCBD | 2 | Irrigated | 7 | 7 | 5 | 0.2 |  | X |  |  |  |  |

Experiments were conducted under conventional tillage with no nutrient limitations; weeds, pests and fungal diseases were chemically controlled. As described in Table 1, experiments were grouped into Sets I, II and III. In Set I, a biparental population of 90 recombinant inbred lines (RIL) derived from the cross between ‘Avalon’ and ‘Glupro’ was evaluated in experiments BAG15 and BAG16. ‘Avalon’ is a cultivar from the UK with high protein content, whereas ‘Glupro’ is a cultivar from the USA with the introgression of the *GPC-B1* gene (Uauy et al., 2006) for high protein content. In Set II, a biparental population of 146 RIL derived from the cross between ‘Baguette 10’ and ‘Klein Chajá’ (Alonso et al., 2018b; Martino et al., 2015; Mirabella et al., 2016) was evaluated in experiments BFE13, BFE14, BFE15 and MJFE17. Both parents are Argentinean spring bread wheat cultivars of good agronomic performance, with contrasting SF and yield components. In Set III, breeding lines derived from crosses between cultivars of diverse origin (CIMMYT, Europe, Argentina), advanced through a modified bulk selection scheme and selected for grain yield and agronomic performance in a wheat breeding program (INTA Balcarce Experimental Station), were evaluated in experiments BLE16 and BLE17 (28 lines), and BLP17 and BLE18 (36 lines).

### Data collection

Grain yield was determined by mechanical harvest of the whole plot (when three-row plots were used) or its five central rows (when seven-row plots were used; Table 1). In the former case, the plot (and harvested) area was 3.5 m^2^, with an inter-plot distance of 0.3 m; in the latter case, the harvested area was 5 m^2^. Grain moisture was obtained from a grain subsample of each plot to calculate dry grain yield (GY; g/m^2^). Grain weight (GW; mg) was determined by counting a 1000-grain sample with an electronic counter, weighing it and dividing total weight by 1000. Grain number per square meter was then calculated as the quotient between GY and GW × 1000.

Spike fertility index was determined at maturity on a sample of 15-20 spikes drawn at random from the harvestable area of each plot prior to mechanical harvest. Spikes were cut at the lowest spikelet level, counted, weighed, and threshed. Spike chaff dry weight (g) was calculated as the difference between total spike dry weight (i.e. before threshing) and total grain weight. Grains were counted using an electronic counter. Then, SF (grains/g) was calculated as the quotient between the number of grains and spike chaff dry weight per sample (Abbate et al., 2013).

Test weight (TW) was measured using a Schopper chondrometer on a 250 g grain sample and expressed as kg/hl. Grain protein content (GPC; %) was determined indirectly by NIRS using a TEC NIR 256 Inlab NIR in 100 g grain samples, previously cleaned and dried at 60°C to constant weight.

### Trait responses after simulated selection

Data from three sets of experiments (Table 1) were used to evaluate responses in GY and other traits after simulated selection for high SF. At each set, the 10% top lines with the highest SF values were selected, based on SF data generated in experiments BAG16 (Set I), BFE13, BFE14, BFE15 and MJFE17 (Set II) and BLE16, BLE17, BLP17 and BLE18 (Set III). This means that, at each set, SF was assessed in only *some* experiments (Table 1). However, the performance of those selected lines and the simulated response after selection (i.e., their differential performance compared with that of the population mean) was evaluated based on data generated in *all* experiments within each set. This also included the experiments in which SF itself was determined, as an independent spike sample was drawn before harvest for SF determination, whereas the response variables were evaluated at harvest. In order to compare the effects of selection for high SF with those for high GY *per se*, a similar approach was followed by carrying out simulated selection for high GY *per se* (10% intensity) and assessing trait responses in all experiments within each set. However, as there is an obvious bias in assessing GY response after selection for GY *per se* in the same experiment, these cases were excluded from the analysis.

Responses after selection for SF or GY *per se* were calculated for GY and other traits as the difference between the mean values of the ‘selected group’ and the general mean of the population at each response experiment. They were then expressed as percentage of the population mean. For each set of experiments, an overall response mean was also calculated.

### Statistical analysis

Grain yield was analyzed with a linear fixed effects model including experiments and genotypes and genotype by experiment interaction effects nested in sets of experiments. Welch’s t-test was used to compare trait responses (in %) after the two simulated selection strategies (i.e., for high SF vs. for high GY *per se*). For this comparison, only those cases that had data for side by side comparisons were used.

## RESULTS

Experiments included in Sets I to III were carried out under different environmental conditions (i.e. different locations, years, growing conditions, plot size, etc.; Table 1). In addition, conspicuous environmental differences between crop seasons due to temperature, radiation and rainfall were observed (Fig. 1). This resulted in significant differences in average grain yield between experiments (p<0.0001); GY differences between genotypes (p<0.0001) were also evident across experiments (Tables 2 and 3). As a consequence of the observed environmental differences and the genetic constitution of the populations under study, conspicuous phenotypic variation was also observed in the remaining traits (Table 3). When analyzed by set, very different population means for all the variables were observed across experiments in each of the three sets, reflecting environmental variation between years and/or locations (Table 3). A summary of results of simulated selection for high SF carried out in each of the three sets is presented in Table 4 (for responses in GY and GN) and Table S1 (for responses in GW, TW and GPC). Selection of the top 10% lines with the highest SF in experiments BAG16 (Set I), BFE13, BFE14, BFE15 and MJFE17 (Set II) and BLE16 and BLE17, BLP17 and BLE18 separately (Set III) always increased GN, and also increased GY except for a few cases of slight decrease in Set III. For each set of experiments, the overall mean response (%) in GY and GN after simulated selection for SF is shown in Table 4. Only for set III the overall mean was negative for GY (−2.0%), while it was positive for GN (8.6%). Across the three set of experiments, GY and GN respectively increased by 5.6 and 13.4% as a result of simulated selection for high SF (Table 4), as compared with the population means. On the other hand, selection responses after simulated selection by high GY *per se* ranged between −9.3% and 46.0% in GY and between −6.8% and 51.2% in GN, considering all experiments. As occurred with simulated selection for high SF, responses after selection for high GY *per se* were positive for all cases except for a few in set III (Table 4).

**Figure 1.**
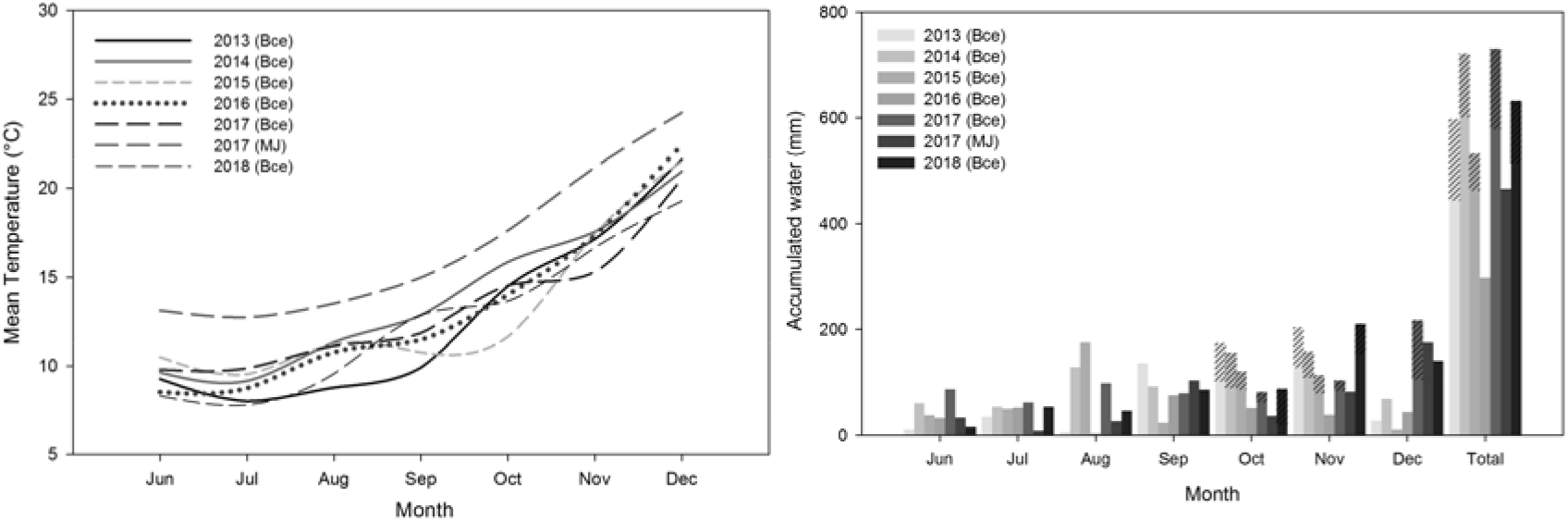
Mean temperature (left) and accumulated rainfall plus irrigation (right) during the June-December period (wheat growing season) at Balcarce (Bce) in 2013 to 2018 and in Marcos Juárez (MJ) in 2017. Stripped bars show accumulated water by irrigation.

**Table 2.** Analysis of variance for grain yield (g/m^2^) across nine experiments. All experiments from Sets I to III (Table 1) were included in the analysis except for MJFE17, which plot size (0.9 m^2^) was deemed too small for accurate grain yield determination.

| Source of variation | DF | Mean Square | F | P-value |
| --- | --- | --- | --- | --- |
| Experiment | 8 | 5155433 | 530.03 | <0.0001 |
| Genotype (set) | 236 | 33324 | 3.43 | <0.0001 |
| Genotype*experiment (set) | 499 | 19126 | 1.97 | <0.0001 |
| Error | 693 | 9727 |  |  |

**Table 3.** Mean (x◻) and standard deviation (S) of spike fertility index (SF, grains/g), grain yield (GY, g/m^2^), grain number per unit area (GN, grains/m^2^), grain weight (GW, mg), test weight (TW, kg/hl) and grain protein content (GPC, %) of I: Avalon/Glupro RIL population (N=90, experiments BAG15 and BAG16); II: Baguette 10/Klein Chajá RIL population (N=146, experiments BFE13, BFE14, BFE15 and MJFE17) and III: advanced breeding lines (N=28, experiments BLE16 and BLE17; N=36, experiments BLP17 and BLE18).

| Set of Experiments | Experiment | SF |  | GY |  | GN |  | GW |  | TW |  | GPC |  |
| --- | --- | --- | --- | --- | --- | --- | --- | --- | --- | --- | --- | --- | --- |
| | | $\bar{x}$ | S | $\bar{x}$ | S | $\bar{x}$ | S | $\bar{x}$ | S | $\bar{x}$ | S | $\bar{x}$ | S |
| I | BAG15 |  |  | 515.9 | 125.0 | 12990 | 3254 | 39.9 | 3.2 | 79.9 | 2.4 | 16.0 | 1.2 |
|  | BAG16 | 70.5 | 11.2 | 314.1 | 81.9 | 9788 | 2554 | 32.3 | 3.4 | 77.1 | 2.1 | 14.5 | 0.7 |
| II | BFE13 | 98.3 | 9.2 | 717.0 | 185.9 | 21960 | 6283 | 33.0 | 4.2 | 77.8 | 11.5 |  |  |
|  | BFE14 | 91.9 | 11.7 | 367.1 | 89.4 | 9964 | 2687 | 37.4 | 4.6 | 73.8 | 2.2 |  |  |
|  | BFE15 | 89.5 | 9.5 | 749.1 | 123.5 | 17824 | 3457 | 42.5 | 3.8 | 80.3 | 1.8 |  |  |
|  | MJFE17 | 101.7 | 12.1 | 432.5 | 134.9 |  |  | 32.4 | 4.7 |  |  |  |  |
| III | BLE16 | 81.5 | 10.2 | 498.9 | 108.4 | 14755 | 3004 | 34.2 | 5.3 |  |  |  |  |
|  | BLE17 | 83.9 | 9.4 | 448.2 | 84.9 | 11302 | 2139 | 39.7 | 4.9 |  |  |  |  |
|  | BLP17 | 86.6 | 11.8 | 383.9 | 46.0 | 11015 | 1770 | 35.3 | 4.1 |  |  |  |  |
|  | BLE18 |  |  | 618.1 | 193.6 |  |  |  |  |  |  |  |  |

**Table 4.**
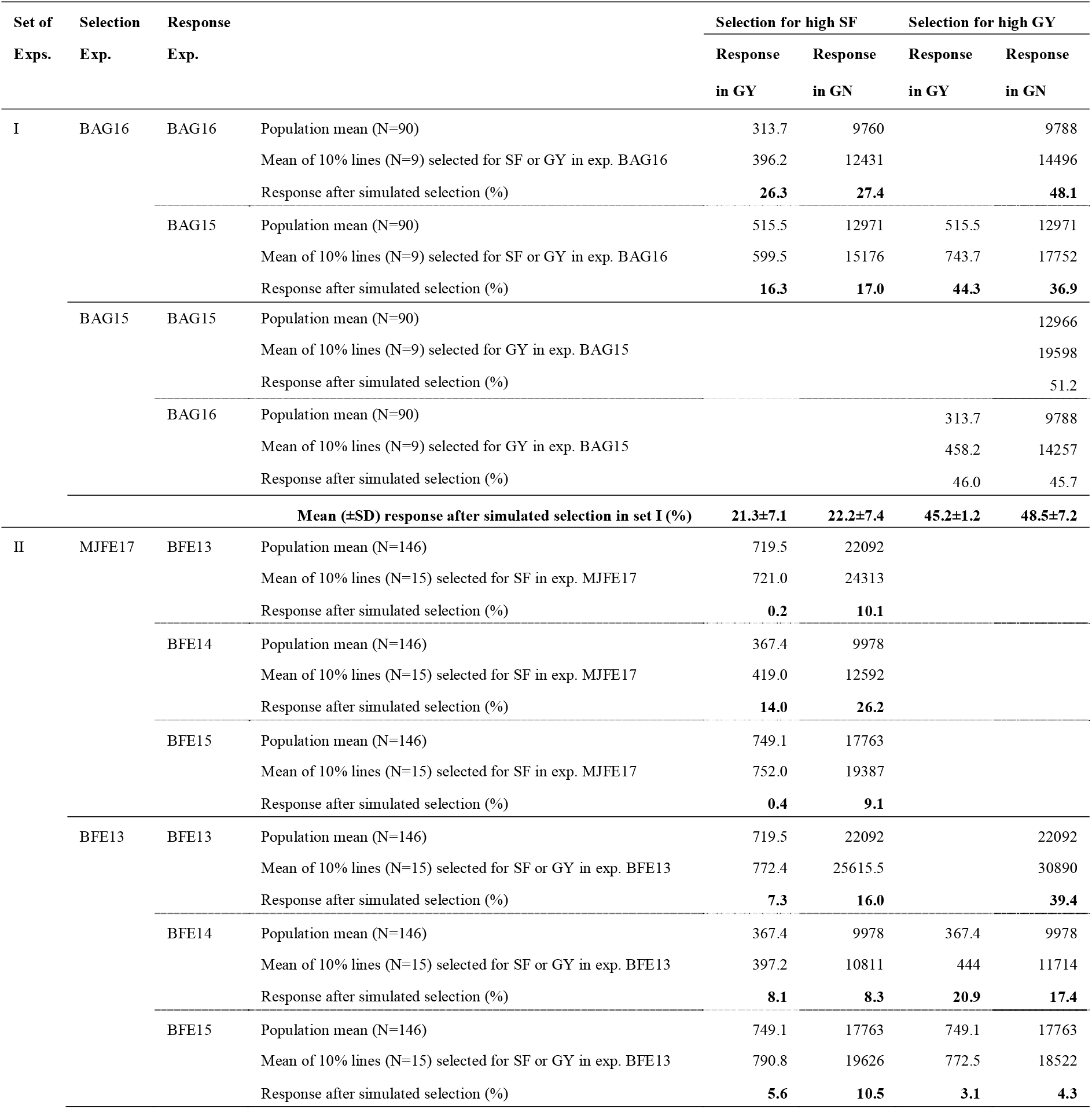
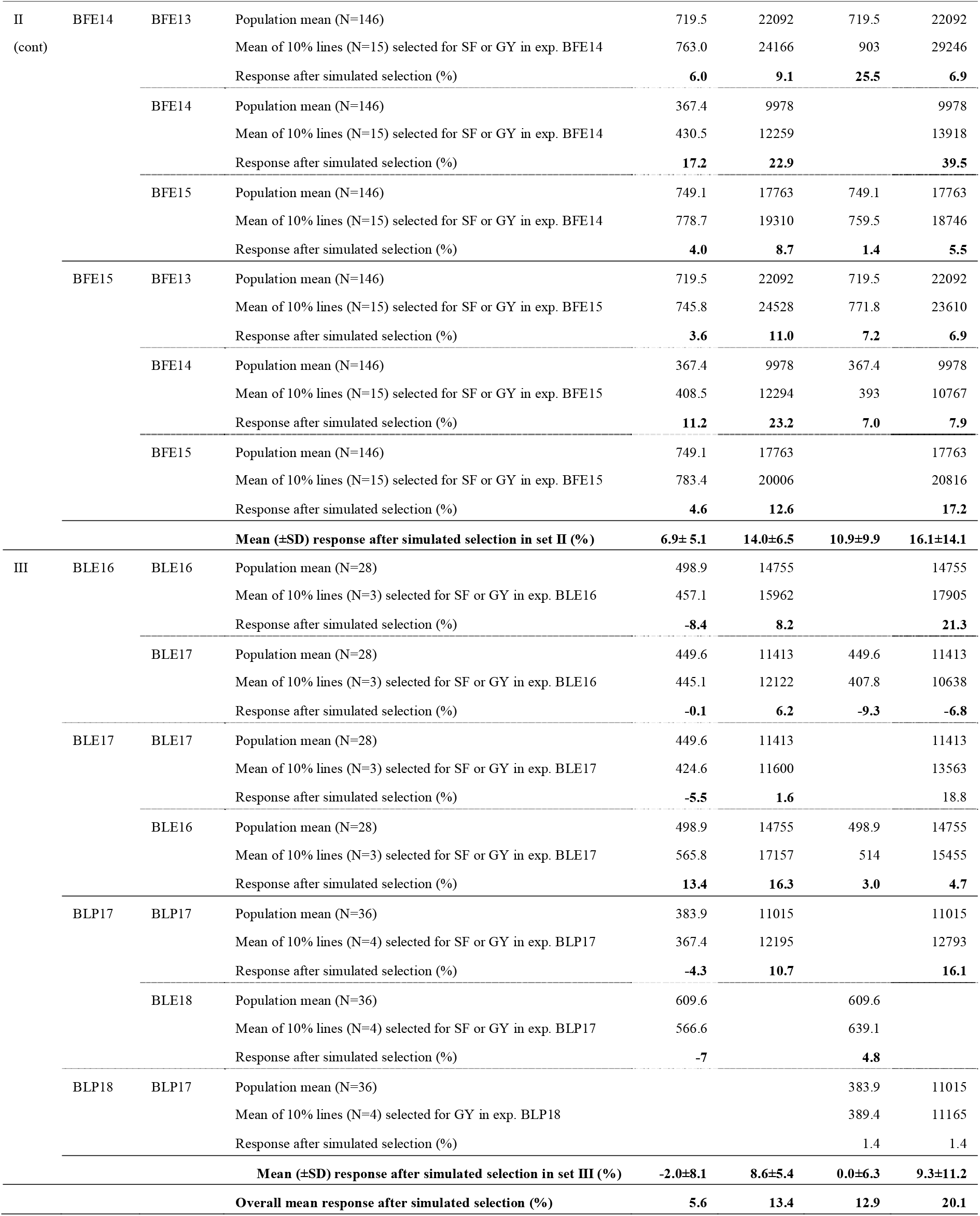
Response after simulated selection for high spike fertility index (SF) and grain yield (GY) per se (i.e., selection of the top 10% lines with the highest SF or GY values at each of three sets of experiments (exps.) carried out between 2013 and 2018 in Balcarce and Marcos Juárez, Argentina) in grain yield (GY; g/m^2^) and grain number per unit area (GN; grains/m^2^). Grain yield responses after selection for GY per se in the same experiment are not shown, as they were not included in the analysis (see Materials and Methods).

Both simulated selection strategies (i.e., by high SF and by high GY *per se*) were compared in terms of the average response attained in GN and GY (Table 5). As a result, both yielded substantially similar average responses (p>0.05) in GN and GY, although the variance was several fold higher when using GY *per se* as the selection criterion for responses in GN.

**Table 5.** T test for comparison of the average response in grain number per unit area (GN; grains/m^2^) and in grain yield (GY; g/m^2^) attained after selection for high spike fertility index (SF) vs. for high GY per se (data extracted from Table 4; only those cases with available data for side by side comparisons were included).

| Variable | Selection criterion | N | Trait response after selection |  | DF | T | P-value |
| --- | --- | --- | --- | --- | --- | --- | --- |
|  |  |  | (%) |  |  |  |  |
|  |  |  | Mean | SEM |  |  |  |
| GN | SF | 16 | 13.1 | 1.73 | 21 | -1.09 | 0.29 |
|  | GY *per se* | 16 | 17.8 | 3.9 |  |  |  |
| GY | SF | 10 | 11.5 | 5.8 | 17 | 0.09 | 0.92 |
|  | GY *per se* | 10 | 11.8 | 4.9 |  |  |  |

As for other yield-related traits, simulated selection for high SF consistently lowered GW (overall decrease of 5.3%; Table S1). As for TW, response after selection for high SF was slightly positive in Set I and slightly negative in Set II, but not significant. Grain protein content, only determined in Set I, was decreased by 6.9% as a result of selection for high SF. On the other hand, average responses after simulated selection for high GY *per se* were positive (Table S1).

## DISCUSSION

Grain yield increase is of paramount importance in every bread wheat breeding program. However, annual increase rates have been steadily declining and would hardly meet the present and future demand. In a recent review, Fischer and Rebetzke (2018) proposed focusing on a number of breeding targets for accelerating grain yield genetic gains, among which they included fruiting efficiency/spike fertility index (SF). When evaluating several selection strategies under different environmental conditions, Alonso et al. (2018b) found that simulated selection for high SF, either solely or along with selection for high grain yield, always resulted in higher and more stable yields than if selecting for high yield alone. In the present study, we show that, regardless of the genetic material or the environmental conditions, selection for high SF alone virtually always increases grain yield in bread wheat.

Independently of the set of experiments, selection for high SF always resulted in GN increases (between 1.6 and 27.4%) and in GY increases in most cases, except for a few in Set III (Table 4). These latter cases, however, occurred in a population of advanced breeding lines, which underwent selection in previous generations as opposed to the RIL populations used in sets I and II. Also, this was the population with the least number of individuals (N=28/36; Table 1). The top 10% lines with the highest SF values amounted to only 3-4, which could have increased sampling error. Nevertheless, overall results suggest that one can select lines with high SF on the basis of data collected from a very small spike sample and sometimes in a different environment than the one in which yield is determined (even including lower yield potential locations as it is Marcos Juárez as compared with Balcarce, in this study) (Menéndez & Satorre, 2007) and expect, at the minimum, a meager increase in yield and/or GN.

Regarding the substantial variation in the magnitude of the responses to selection observed across sets and experiments within sets (Table 3), several possible underlying causes could be considered. For instance, variations might have occurred due to unmeasured sampling errors, sample size, and different genetic and genetic by environment interaction effects of the traits under evaluation. In this sense, additional data sets comprising a wider spectrum of germplasm and environments/years of evaluation should be incorporated in the analysis in order to reduce these errors. On the other hand, the heritability of each trait as well as the degree of association between the trait used for selection and the response trait might have been other sources of variation in the observed responses to selection. For instance, in the case of the RIL population of Set II, the heritability values of the traits (*h^2^*_SF_=0.84; *h^2^*_GY_=0.28; *h^2^*_GN_=0.35; Alonso et al. 2018b) and the correlations between the traits, both genetic (r_SF-GY_=0.36, r_SF-GN_=0.55; Alonso et al. 2018b) and phenotypic (r_SF-GY_=0.29, r_SF-GN_=0.44; Table S2), provide an idea of how these relationships could affect the responses. Spike fertility index, used here as a selection criterion, has high heritability with low environmental influence, which makes it fairly stable across successive generations. The trait response after such selection is expected to be positively correlated with the degree of association between the selection and evaluation traits; indeed, in this study, the response was better in GN than in GY. In turn, the year-to-year variations in the response of these traits are in line with their heritabilities (low to moderate) and with their great sensitivity to environmental effects. Therefore, selecting for high SF would give stability to the response in GN and GY, whereas these latter traits’ low/moderate values of both heritability and genetic/phenotypic correlation with SF would increase variability in such response.

Some trade-offs were observed: in general, GW, GPC and TW tended to decrease with selection for high SF. Nevertheless, genotypes with all high SF, GY, GW, GPC and TW were detected (data not shown); therefore, these trade-offs could be avoided by concurrent selection for all traits. Furthermore, all three populations used in the present study had a relatively low number of individuals if considering the scale of an actual breeding program. Provided that the prerequisite of existence of genetic variability for the traits is met (as it was the case in the present study; Table 3), increasing population numbers is likely to yield even better results when selecting for high SF. This way, while increasing GY and GN, one could concurrently maintain desirable GW, TW and GPC values. Phenotypic correlations between variables were generally positive (Table S2) except for the ones between GN and GW (r=−0.18) and SF and GW (r=−0.2). These low values, however, confirm the idea that the negative relationship between these variables is not complete.

Independently of the genetic constitution of the breeding material and/or the environmental conditions, selections for high SF, as determined in a small spike sample, appear to result in consistently increased grain number per unit area (and yield) in bread wheat. As virtually similar average responses to selection in GN and GY were attained when using SF or GY *per se* as selection criteria (Table 5), selection for SF alone has the potential of increasing the throughput and efficiency of the breeding process, while lowering costs. Spike fertility index determination entails drawing samples in the field, weighing and threshing (which, although cheaper, might be more labor-intensive and time-consuming than GY determination), but it is a process amenable to automation. This could translate into a massive screening of lines for SF in small plots and subsequent multi-environment yield trials that only include a subset of the lines initially screened. This turns SF into a very promising selection target in breeding programs aimed at increasing grain yield. Nevertheless, the effectiveness of carrying out selection for high SF in spaced plants, as it would be the case of early generations of a breeding program, remains yet to be established.

## ACKNOWLEDGEMENTS

We thank members of the Grupo Trigo Balcarce (EEA Balcarce INTA) and Mejoramiento de Trigo y Biotecnología (EEA Marcos Juárez INTA) for help with the experiments and technical assistance. Scholarships granted to M.F. Franco and M.P. Alonso by CONICET and to N.E. Mirabella by INTA are acknowledged. This research was partially funded by INTA (PNCyO 1127044).

## DATA AVAILABILITY

The dataset of the study is available from the authors upon reasonable request.

## CONFLICTS OF INTEREST

The authors declare that they have no conflicts of interest.

## AUTHOR CONTRIBUTIONS

ACP, MPA and NEM designed the study; ACP, MPA, NEM, JSP, MFF, and ML performed experiments at Balcarce and LSV performed the experiment at Marcos Juárez; MPA, NEM and JSP processed SF samples; NEM, MPA and ACP analyzed the data; MFF prepared all figures of the manuscript; ACP wrote the paper with input from all authors.

---

**Table S1.**
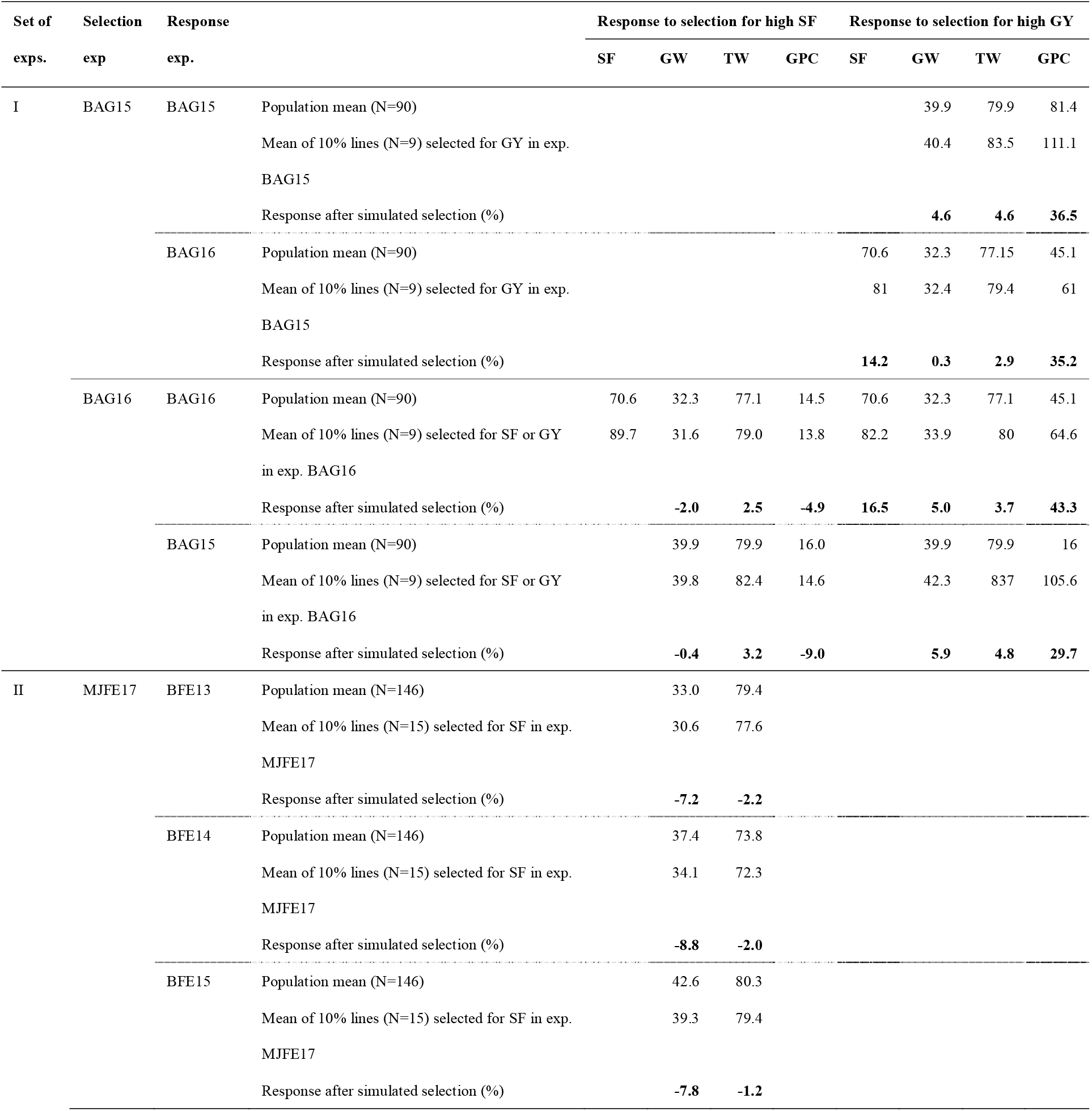
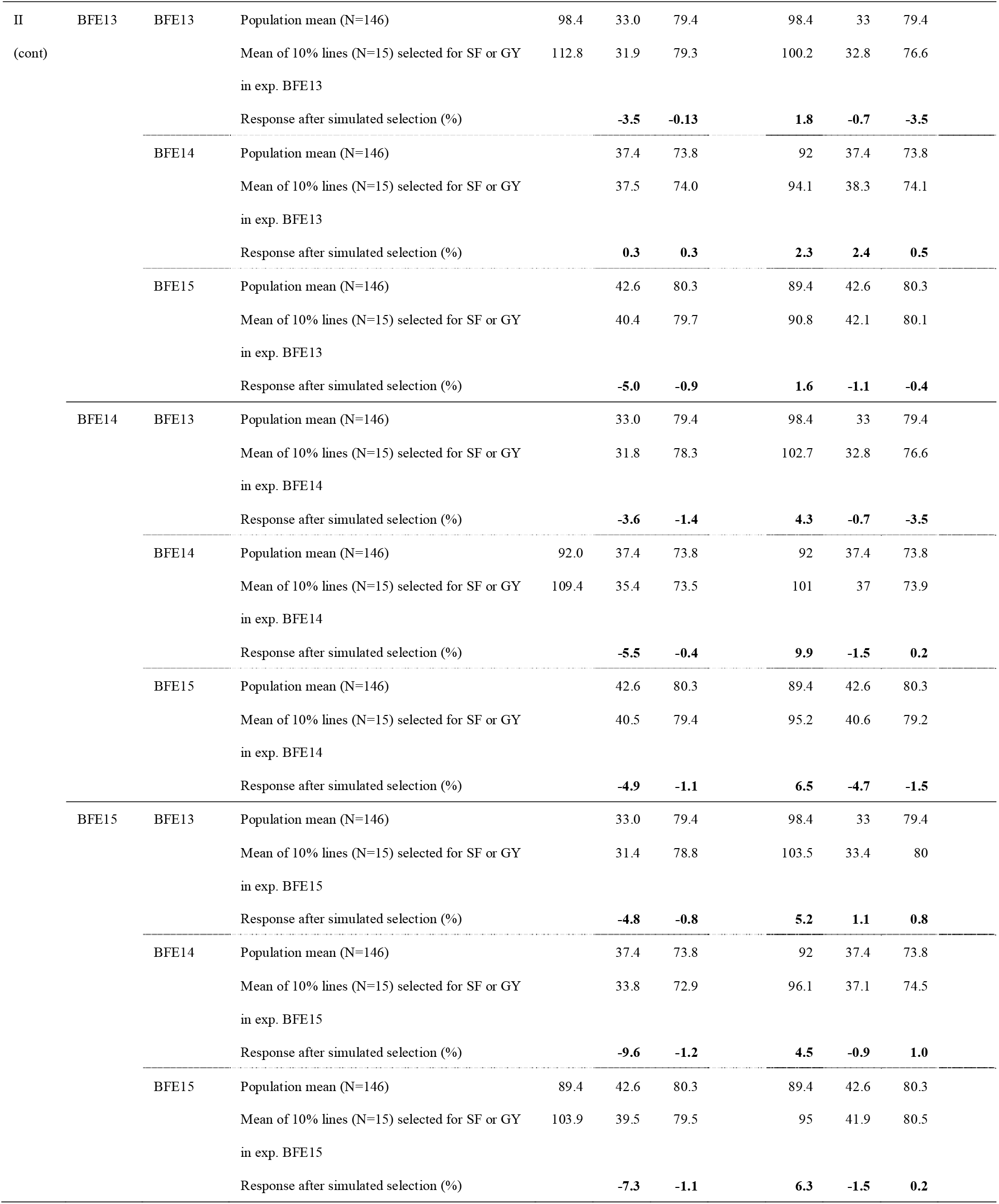
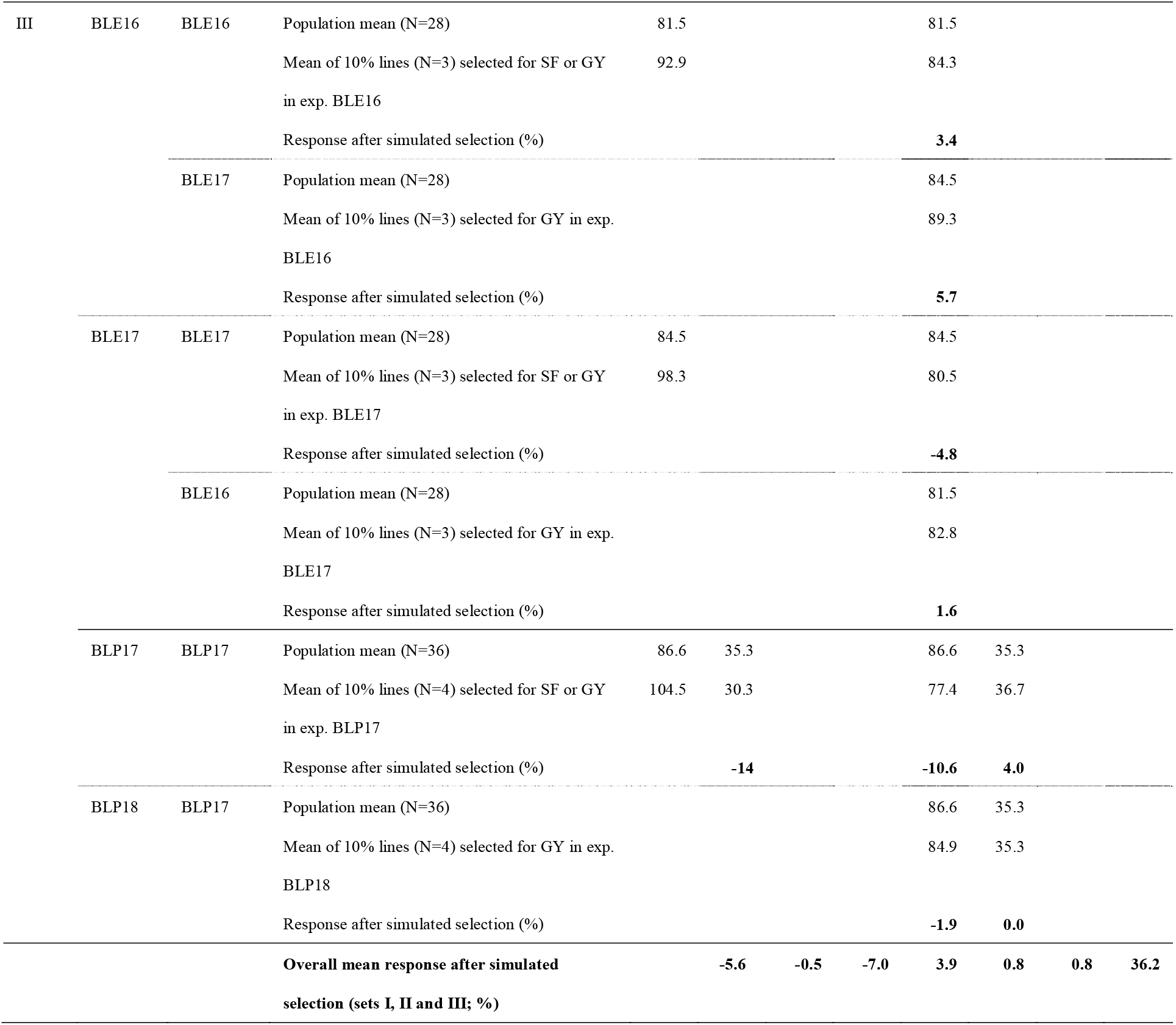
Response to simulated selection for high spike fertility (SF) or grain yield (GY) per se (i.e., selection of the top 10% lines with the highest SF or GY values at each of three sets of experiments (exps.) carried out between 2013 and 2018 in Balcarce and Marcos Juárez, Argentina) in spike fertility index (SF; grains/g), grain weight (GW; mg), test weight (TW; kg/hl) and grain protein content (GPC; %).

**Table S2.** Pearson correlation coefficients between spike fertility index (SF, grains/g), grain yield (GY, g/m^2^), grain number per unit area (GN, grains/m^2^), grain weight (GW, mg) and test weight (TW, kg/hl), calculated with data from sets of experiments I, II and III. All coefficients are significant (p<0.05) except where noted (ns).

|  | GY | GN | GW | TW |
| --- | --- | --- | --- | --- |
| SF | 0.29 | 0.44 | -0.20 | 0.00<sub>ns</sub> |
| GY |  | 0.91 | 0.24 | 0.28 |
| GN |  |  | -0.18 | 0.19 |
| GW |  |  |  | 0.21 |

## Notes

### Competing Interest Statement

The authors have declared no competing interest.

## REFERENCES

Abbate, P., Andrade, F., Lazaro, L., Bariffi, J., Berardocco, H., Inza, V., & Marturano, F. (1998). Grain yield increase in recent Argentine wheat cultivars. Crop Science, 38, 1203–1209. doi: 10.2135/cropsci1998.0011183X003800050015x.

Abbate, P. E., Pontaroli, A. C., Lázaro, L., & Gutheim, F. (2013). A method of screening for spike fertility in wheat. The Journal of Agricultural Science, 151, 322–330. doi: 10.1017/s0021859612000068.

Acreche, M. M., Briceño-Félix, G., Sánchez, J. A. M., & Slafer, G. A. (2008). Physiological bases of genetic gains in Mediterranean bread wheat yield in Spain. European journal of agronomy, 28, 162–170. doi: 10.1016/j.eja.2007.07.001.

Alonso, M.P., Abbate, P.E., Mirabella, N. E., Aramburu Merlos, F., Panelo, J.S., & Pontaroli, A.C. (2018a). Analysis of sink/source relations in bread wheat recombinant inbred lines and commercial cultivars under a high yield potential environment. European Journal of Agronomy, 93:8–87. doi.org/10.1016/j.eja.2017.11.007

Alonso, M., Mirabella, N., Panelo, J., Cendoya, M., & Pontaroli, A. (2018b). Selection for high spike fertility index increases genetic progress in grain yield and stability in bread wheat. Euphytica, 214, 112. doi: 0.1007/s10681-018-2193-4.

Borrás, L., Slafer, G. A., & Otegui, M. E. (2004). Seed dry weight response to source–sink manipulations in wheat, maize and soybean: a quantitative reappraisal. Field Crops Research, 86, 131–146. doi: 10.1016/j.fcr.2003.08.002.

CIMMYT. (2019). Retrieved Accessed 2019 Sept 15, from https://www.cimmyt.org/news/food-security/

Elía, M., Savin, R., & Slafer, G. A. (2016). Fruiting efficiency in wheat: physiological aspects and genetic variation among modern cultivars. Field Crops Research, 191, 83–90. doi: 10.1016/j.fcr.2016.02.019.

Ferrante, A., Cartelle, J., Savin, R., & Slafer, G. A. (2017). Yield determination, interplay between major components and yield stability in a traditional and a contemporary wheat across a wide range of environments. Field Crops Research, 203, 114–127. doi: 10.1016/j.fcr.2016.12.028.

Ferrante, A., Savin, R., & Slafer, G. A. (2012). Differences in yield physiology between modern, well adapted durum wheat cultivars grown under contrasting conditions. Field Crops Research, 136, 52–64. doi: 10.1016/j.fcr.2012.07.015.

Fischer, R. (2007). Understanding the physiological basis of yield potential in wheat. The Journal of Agricultural Science, 145, 99. doi: 10.1017/S0021859607006843.

Fischer, R. (2011). Wheat physiology: a review of recent developments. Crop and Pasture Science, 62, 95–114. doi: 10.1071/CP10344.

Fischer, R., & Rebetzke, G. (2018). Indirect selection for potential yield in early-generation, spaced plantings of wheat and other small-grain cereals: a review. Crop and Pasture Science, 69, 439–459. doi: 10.1071/CP17409.

Fischer, R. (1984). Wheat. In W. Smith & S. Banks (Eds.), Proceedings of symposium on potential productivity of field crops under different environments (pp. 129–154). IRRI, Los Baños, Philippines.

Foulkes, J., Rivera, C., Trujillo, E., Sylvester-Bradley, R., & Reynolds, M. (2015, March 24-26). Achieving a step-change in harvest index in high biomass wheat cultivars. Paper presented at the Proceedings of the International TRIGO (Wheat) Yield Potential Workshop 2015, Obregón, Sonora, Mexico.

Foulkes, M. J., Slafer, G. A., Davies, W. J., Berry, P. M., Sylvester-Bradley, R., Martre, P., … Reynolds, M. P. (2010). Raising yield potential of wheat. III. Optimizing partitioning to grain while maintaining lodging resistance. Journal of Experimental Botany, 62, 469–486. doi: 10.1093/jxb/erq300.

García, G. A., Serrago, R. A., González, F. G., Slafer, G. A., Reynolds, M. P., & Miralles, D. J. (2014). Wheat grain number: identification of favourable physiological traits in an elite doubled-haploid population. Field Crops Research, 168, 126–134. doi: 10.1016/j.fcr.2014.07.018.

González, F., Terrile, I. I., & Falcón, M. (2011). Spike fertility and duration of stem elongation as promising traits to improve potential grain number (and yield): variation in modern Argentinean wheats. Crop Science, 51, 1693–1702. doi: 10.2135/cropsci2010.08.0447.

Lázaro, L., & Abbate, P. E. (2012). Cultivar effects on relationship between grain number and photothermal quotient or spike dry weight in wheat. The Journal of Agricultural Science, 150, 442–459. doi: 10.1017/s0021859611000736.

Lynch, J. P., Doyle, D., McAuley, S., McHardy, F., Danneels, Q., Black, L. C., … Spink, J. (2017). The impact of variation in grain number and individual grain weight on winter wheat yield in the high yield potential environment of Ireland. European journal of agronomy, 87, 40–49. doi: 10.1016/j.eja.2017.05.001.

Martino, D. L., Abbate, P. E., Cendoya, M. G., Gutheim, F., Mirabella, N. E., & Pontaroli, A. C. (2015). Wheat spike fertility: inheritance and relationship with spike yield components in early generations. Plant Breeding, 134, 264–270. doi: 10.1111/pbr.1226.

Menéndez, F. J., & Satorre, E. H. (2007). Evaluating wheat yield potential determination in the Argentine Pampas. Agricultural systems, 95, 1–10. doi: 10.1016/j.agsy.2007.03.004.

Mirabella, N., Abbate, P., Ramirez, I., & Pontaroli, A. (2016). Genetic variation for wheat spike fertility in cultivars and early breeding materials. The Journal of Agricultural Science, 154, 13–22. doi: 10.1017/S0021859614001245.

Shearman, V., Sylvester-Bradley, R., Scott, R., & Foulkes, M. (2005). Physiological processes associated with wheat yield progress in the UK. Crop Science, 45, 175–185. doi: 10.2135/cropsci2005.0175.

Slafer, G. A., Andrade, F. H., & Satorre, E. H. (1990). Genetic-improvement effects on pre-anthesis physiological attributes related to wheat grain-yield. Field Crops Research, 23, 255–263. doi: 10.1016/0378-4290(90)90058-J.

Slafer, G. A., Elia, M., Savin, R., García, G. A., Terrile, I. I., Ferrante, A., … González, F. G. (2015). Fruiting efficiency: an alternative trait to further rise wheat yield. Food and Energy Security, 4, 92–109. doi: 10.1002/fes3.59.

Terrile, I. I., Miralles, D. J., & González, F. G. (2017). Fruiting efficiency in wheat (Triticum aestivum L): Trait response to different growing conditions and its relation to spike dry weight at anthesis and grain weight at harvest. Field Crops Research, 201, 86–96. doi: 10.1016/j.fcr.2016.09.026.

Uauy, C., Distelfeld, A., Fahima, T., Blechl, A., & Dubcovsky, J. (2006). A NAC gene regulating senescence improves grain protein, zinc, and iron content in wheat. Science, 314(5803), 1298–1301.

